# The balance between ATR and DDK activities controls TopBP1-mediated locking of dormant origins at the pre-IC stage

**DOI:** 10.1101/2023.11.29.569233

**Authors:** Stéphane Koundrioukoff, Su-Jung Kim, Nathan Alary, Antoine Toffano, Rodrigo Melendez-Garcia, Xia Wu, Yaqun Liu, Stefano Gnan, Sami El-Hilali, Olivier Brison, Filippo Rosselli, Chun-Long Chen, Michelle Debatisse

## Abstract

Replication stress, a major hallmark of cancers, and ensuing genome instability source from impaired progression of replication forks. The first line of defense against fork slowing is compensation, a long-described process that elicits firing of otherwise dormant origins. It remains unclear whether compensation requires activation of the DNA replication checkpoint or passively results from lengthening of the window of time during which dormant origins can fire when fork progression slows, or both. Using molecular DNA combing we show here that a linear relationship ties inter-origin distances to fork speeds, independently of the checkpoint status. We called this line “stressline” and further show that its slope enables precise quantification of the compensation efficiency. Comparison of the slopes in different genetic backgrounds reveals that compensation requires ATR, not CHK1, while TopBP1 and CDC7/DBF4 repress dormant origin activation. These results strongly suggest that TopBP1 locks dormant origins at the pre-IC stage and that ATR and DDK oppose to control the conversion of dormant pre-ICs into functional salvage origins. Both passive and active processes thus contribute to compensation. Moreover, Repli-seq and OK-seq analyses confirm the activating role of ATR and permit development of ATRAP-seq, a new procedure allowing mapping of early constitutive origins.

## Introduction

Duplication of large metazoan genomes relies on the firing of thousands of replication origins that must be tightly coordinated to avoid under- or over-replication and their deleterious consequences on genome stability^1^. Origin building is a step-wise process, which provides multiple opportunities for regulation. The first step is the ORC-directed recruitment of two MCM hexamers in head-to-head orientation, constituting the pre-replication complex (pre-RC)^2,3^. This process is well conserved across eukaryotes although the modalities of ORC recruitment itself are not^4^. The second step occurs when the CDC7-DBF4 kinase (DDK) and S phase cyclin-dependent kinases (CDKs) phosphorylate the MCMs, eliciting recruitment of DUE B^5^, the Treslin-MTBP complex, TopBP1, CDC45^2^, and the GINS-DONSON complex^6–8^ to form the pre-IC. The third step consists in evicting DUE B, Treslin-MTBP and TopBP1, permitting emergence of the replicative helicase (CDC45/MCMs/GINs: CMG complex) associated with Pol ε (CMGE complex)^9–11^. The order in which and how these proteins are recruited, then released for some of them, as well as their exact function(s) in CMGE setting remains incompletely documented, particularly in mammalian cells. A fourth round of protein recruitment finally sets functional replisomes that initiate replication according to a cell type-specific timing program^12^. This program is regulated at the level of large-scale domains^13^ by a complex combination of cis-regulatory sequences^14^, trans-acting factors and 3D genome architecture^15–17^.

Upon stress, many more origins fire along timing domains engaged in the replication process than in a normal S phase, which sustains local replication rate^18^. It has long been proposed that extra-origins source from the pre-RC pool, which greatly exceeds the number of origins activated during a normal S phase^19^. However, many questions remain about the mechanisms preventing supernumerary pre-RC to give rise to functional origins in unstressed cells. Notably, the stage at which the building process of dormant origins is arrested, and whether their activation and/or completion under stress is an active or a passive process. The passive hypothesis proposes that dormant origins are ready to start but inactivated by forks coming from earlier origins of the domain. In this hypothesis, the level of fork slowing dictates the window of time preceding their inactivation, and the density of initiation events in consequence. Alternatively, or in addition, dormant origins may be halted at an intermediate step of the building process. In this case, the DNA damage checkpoint (DRC)^20^ is an obvious candidate for triggering their completion and activation. However, it has long been established that activation of the ATR-CHK1 pathway delays replication of domains that have not yet begun to replicate, thereby preventing escalation of fork slowing. It therefore remains to explain how the same pathway could simultaneously activate dormant origins locally and repress initiation globally^21^.

A possible explanation for these opposite roles of the DRC comes from previous work showing that treatments with low concentrations of aphidicolin (Aph), an inhibitor of the replicative DNA polymerases, trigger a loose DRC response characterized by loading of ATR on the chromatin but no, or weak, CHK1 activation^22^. A rationale for these observations recently came from single-molecule experiments showing that ATR constantly monitors the level of RPA at the forks under low stress conditions. The accumulation of RPA at severely impeded forks under more stringent stress promotes ATR crowding, which triggers signal amplification and a shift from loose to full DRC activation^23^. Together, the two sets of results therefore suggest that ATR, by itself, promotes local events while CHK1 induces additional remote responses according to stress stringency.

In addition, TopBP1, the major ATR activator during S phase^20^, Treslin-MTBP^24^ and DONSON^6–8^ play important role in both cell cycle checkpoints and CMGE setting, which favors crosstalk between helicase activity and cell cycle regulation. TopBP1 (Dpb11/Rad4/Xmus101) is essential for pre-IC building in budding and fission yeasts^25^, and in the xenopus egg model^11,26,27^. However, while it is generally accepted that this latter function of the protein is conserved in mammalian cells, several reports show that, at short term, TopBP1 depletion impacts the replication process neither in human somatic cells nor during the development of mouse embryos. Indeed, TopBP1 depletion results in the progressive accumulation of DNA damages across the following cell cycles, then to senescence or apoptosis depending on the experimental models^28–31^. Since the protein caps the pre-IC and is released before firing, as in other eukaryotes, its functions in origin setting and, potentially, in coupling replication initiation with DRC activation therefore remain to be understood.

## Results

### A linear relationship links the inter-origin distances to fork speeds

In order to determine the impact of fork slowing on dormant origin activation, we used molecular DNA combing (Fig. 1A, B and S1A). This single-molecule technique gives access to fork speed and to the distances separating two origins (Inter-Origin Distances: IODs), a parameter that reflects the density of initiation events along individual genomic domains. We treated human cells with 0.03 to 0.3 μM of Aph, which leads to gradual fork slowing (Fig. S1B) and progressive recruitment of ATR on the chromatin, but no CHK1 phosphorylation^22^. These conditions thus avoid the many consequences of CHK1 activation on replication fitness and cell cycle progression^20^ that could superimpose compensation and blur interpretation of the results. In addition, DNA counter-staining permitted us to select DNA molecules longer than 0.5 Mb, which avoids bias toward small IODs resulting from analysis of short molecules^32^.

**Figure 1:**
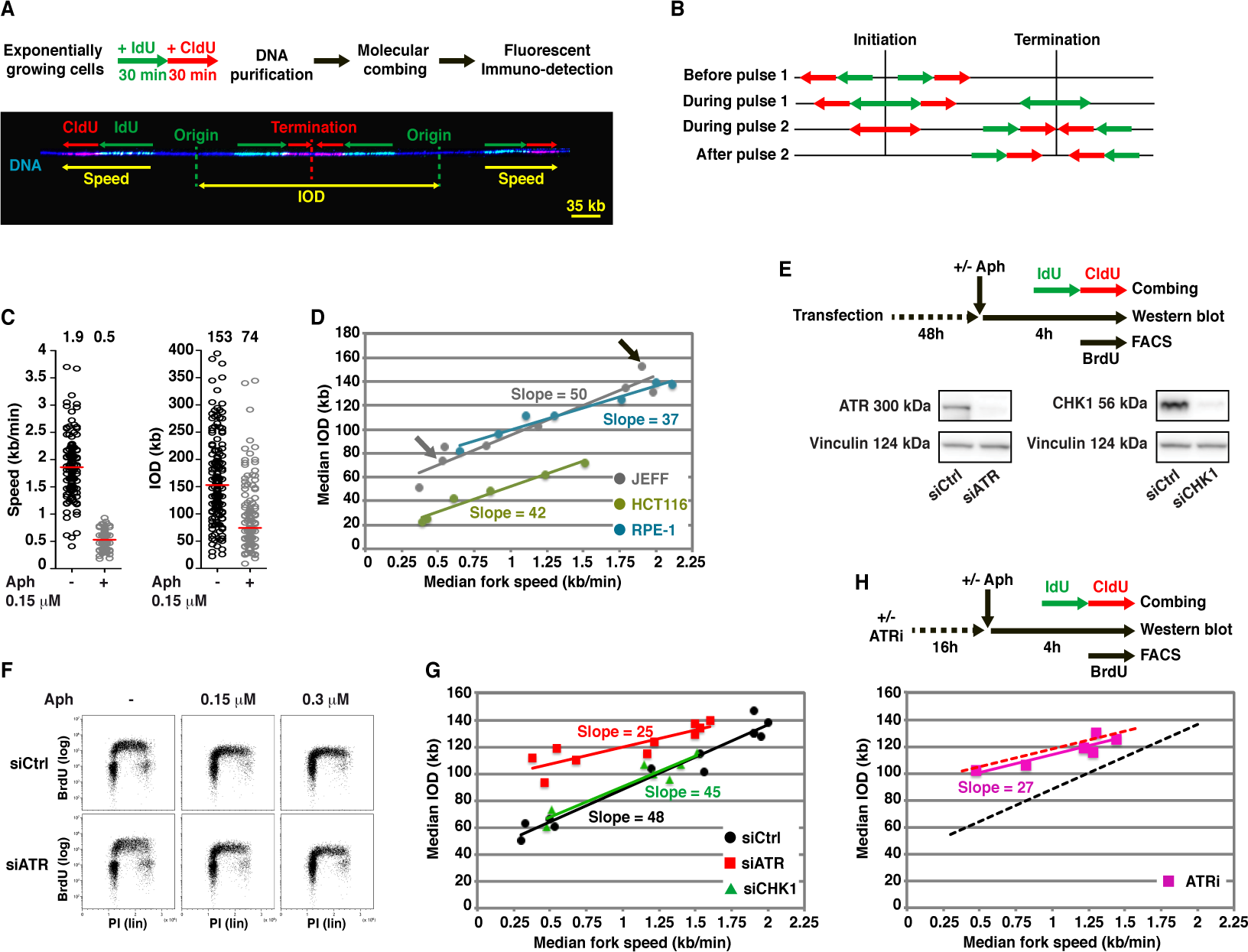
Efficient compensation relies on ATR but not on CHK1. **A:** Upper panel; principle of molecular DNA combing labelling. Pulse-labelling with IdU and CldU (30 min each) allows determination of fork directionality. Lower panel; DNA molecules displaying two initiation events, and schematic representation of signals used to determine fork speeds and inter-origin distances (IOD) (yellow), DNA is counterstained in blue (methods). B: Schematic representation of the patterns observed in cells treated as in A (adapted from32), depending on when the considered event had occurred relative to the pulses. C: Determination of the median fork speed (left), and median IOD (right) in cells treated as indicated (n>150 measures for each parameter, the medians are indicated above the cloud of measures). D: Stresslines obtained with the indicated cell lines (n>150, as in C for each point). Arrows underline the points corresponding to the experiments shown in C (same color-code). E: Upper panel; Experimental procedure for transfection experiments. When added, Aph was present during the last 4 h before recovery (including the labelling periods). Lower panel; western blots showing the efficiency of ATR and CHK1 depletion 48 h after transfection. F: FACS analysis of cell populations treated as in E (BrdU/FACS). The experiment has been done once. G: Stresslines obtained with cells transfected with the indicated siRNAs and treated as in E (combing) (n>150 as in C for each point). H: Upper panel; experimental procedure for cells treated with ATRi. Lower panel; Stressline for ATRi. Dotted lines show the lines obtained with siATR (red) and siCtrl (black) for comparison.

Analysis of lymphocytes immortalized by EBV (JEFF cells) showed that even the lowest levels of fork slowing impact the IODs and, strikingly, that a linear relationship links the median fork speeds to the median IODs (Fig. 1C, D). We also studied hTERT-immortalized epithelial retina cells (RPE-1) and colorectal carcinoma cells (HCT116). Comparison of the results obtained with untreated cells, referred hereafter to the start-point, showed important variations from cell line to cell line, which most likely reflects differences in their genetic background. In contrast, the linear relationship between fork speeds and IODs was maintained in all examined cell lines (Fig. 1D), making this property a constant of the replication process. We called this straight line “stressline”. The linearity may passively result from processive inactivation of dormant origins by forks coming from earlier ones, or may stem from progressive increase in ATR activity upon gradual fork slowing, or both. To address this question, we determined whether the stressline linearity persists in JEFF lymphoblasts in which various DRC proteins have been depleted or inhibited.

### DRC-independent and DRC-dependent processes contribute to compensation

We depleted JEFF cells of ATR, or CHK1 as a control, and first performed fluorescence-activated cell sorting (FACS) analysis to ensure that the cell distribution in the cycle is not, or marginally, perturbed in cells treated as described in Figure 1E. We found that neither ATR (Fig. 1F) nor CHK1 (Fig. S1C) depletion affect this distribution whether the cells were treated or not with Aph. Molecular combing showed that the linear relationship linking fork speeds to IODs persists whether the cells were transfected with siCtrl (control siRNAs, no known target), siATR or siCHK1 (Fig. 1G). We conclude that the linearity is DRC-independent. Analysis of the start-points showed a decrease in the median fork speed without changes in the median IOD in ATR-depleted cells (Fig. 1G). In CHK1 depleted cells, the start-point also displayed a decrease in the median fork speed, accompanied by a shortening of the median IODs (Fig. 1G), which confirms results previously obtained in JEFF cells^33^. Together, our observations thus indicate that the linearity is independent of the start-point and most probably relies on a passive mechanism.

We then focused on the slope of the stressline. Strikingly, ATR-depleted cells displayed a strongly flattened line compared to ATR-proficient cells (Fig. 1D, G), suggesting that far less dormant origins fire for equivalent fork speed reduction in this genetic background. To strengthen this conclusion, we performed statistical analysis of the results. Compared to non-transfected cells, transfection with siCtrl did not impact the slope (50 versus 48, respectively (F(1, 16) = 7.92−10-2, p = 0.78 (ANOVA test; Methods)) (Fig. 1D, G), we therefore combined the two data sets to create a reference stressline (Ref-L). Note that the same statistical test was used below for all comparisons of stressline slopes. We found that the slope resulting from ATR depletion (25) significantly differs from that of Ref-L (F(1, 25) = 11.54, p = 2.2−10^−3^). In contrast, the slope resulting from CHK1 depletion (45) does not (F(1, 23) = 9.29−10^−2^, p = 0.76) (Fig. 1G), indicating as expected that CHK1 is not involved in compensation. The latter conclusion agrees with that of a previous study that used a completely different technique^34^, which validates the stressline approach.

To confirm the role of ATR in dormant origin activation, the cells were treated with 100 nM of the ATR inhibitor VE-822 (ATRi) (Fig. 1H, upper panel), a concentration that strongly reduces ATR-dependent CHK1 phosphorylation upon stringent replication stress (Fig. S1D). FACS analysis of cells treated with ATRi+Aph confirmed that S phase progression is independent of ATR in the conditions used here (Fig. S1E). Molecular combing showed that the linearity persists, and the stresslines slope (27) was similar whether the cells were treated with ATRi or depleted of the protein (F(1, 12) = 1.74−10^−2^, p = 0.89) (Fig. 1G, H). Compared to ATR-depleted cells, the start-point of ATRi-treated cells displays a stronger reduction of the median fork speed and a modest reduction of the median IOD (Fig. 1H), a phenotype reminiscent of that previously described by others^35^. We concluded that the stressline slope, like the linearity, is independent of the start-point (Fig. 1D) and that ATR activity is key to efficient activation of dormant origins that have not yet been passively inactivated. The slope thus reflects the magnitude of the response to fork slowing and provides a unique tool for distinguishing “active compensation” from “passive compensation”.

### Combination of ATRi and Aph reveals discrete origins

The organization of timing domains, a major parameter of the replication dynamics, escapes molecular combing analyses. We therefore used Repli-Seq^36^ (Fig. 2A) to further study the impact of ATR inhibition in cells challenged with low Aph concentrations. We compared the profiles of untreated cells to those of cells treated with ATRi or Aph alone and ATRi+Aph (Fig. 2B and S2A, B). By visual inspection, the profiles in untreated cells and cells treated with ATRi remained unchanged. Global profile was also unchanged in Aph-treated proficient cells, with the exception of a few specific loci^37^. Strikingly, cells treated with ATRi+Aph displayed discrete peaks preferentially localized in early replicating domains (G1/S1) (Fig 2B). We checked the robustness of these data by performing independent biological experiments, focusing on cells sorted in G1/S1 following treatment with ATRi+Aph 0.3 or 0.6 μM (n=3 for each Aph concentration). After normalization (Methods), data obtained from the initial experiment and these replicates gave highly reproducible profiles (Fig 2C and S2A, B). We identified 5407 consensus peaks (peaks detected on the mean profile of all ATRi+Aph samples) within early replicating genome (S50 values > 0.4 on a scale of 0 (earliest) to 1 (latest), the S50 being the moment in S phase when a sequence has been replicated in 50% of the cells (Methods)) (Fig 2C and S2C, left and middle panels). Those peaks, called ATRAP (ATRi+Aphidicolin) peaks, were significantly larger (p < 2.2−10^−16^, one-sided paired t-test) in cells treated with 0.3 μM than with 0.6 μM of Aph (Fig. 2D), which is consistent with the difference in fork slowing in the two conditions. We thus conclude that ATRAP peaks result from the combination of deficiency in active compensation and slow fork progression that together allow visualization of neo-synthesized DNA originating from a pool of strong ATR-independent origins.

**Figure 2:**
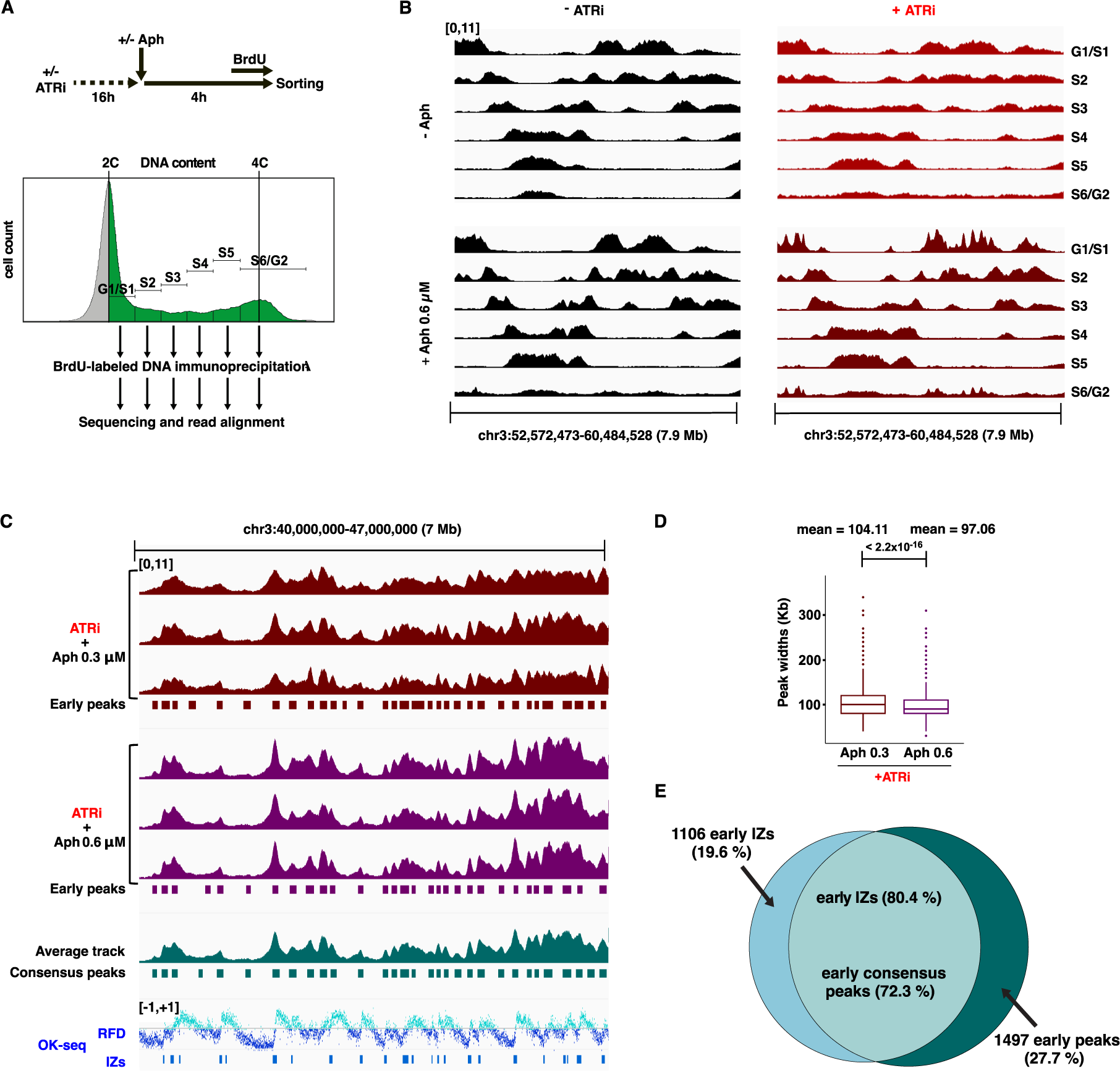
Aph treatment reveals constitutive origins in ATR deficient cells. **A:** Experimental scheme for Repli-seq analyses. Newly synthesized DNA was labeled *in vivo* as described in the upper panel. BrdU labelling: 1h for cells not treated with Aph and 3h for cells treated with Aph to reach sufficient amounts of labelled DNA. Cells were then FACS-sorted in 6 fractions, from G1/S1 to S6/G2M, according to their DNA content. DNA was prepared and treated as indicated (methods). B: Sequencing of BrdU labelled DNA in the indicated growth conditions. The normalized read density profiles (Methods) of each fraction are shown in 50 kb sliding windows (1 kb step) and range between 0 and 11. C: Profiles of G1/S1 fractions (presented as in B). Three biological replicates have been performed for each treatment. The average profile of the 6 replicates (average track) and the corresponding consensus peaks (ATRAP-peaks) are shown. At the bottom, early initiation zones (IZs with replication timing values S50 ≤ 0.4) identified by OK-seq, aligned with Repli-seq data. Replication fork directionality (RFD), (ranging from −1 to +1) is shown. D: Boxplots showing the width distribution of ATRAP-seq peaks. Bounds of box: 25th and 75th percentiles; center line: median; lower (upper) whisker: lower (upper) bound - (+) 1.5 x Interquartile range. Only the one-to-one match peaks (n=4888) in both conditions were included. The p-value was obtained by one-sided paired t-test. E: Venn diagram presenting the overlap between early OK-seq IZs and ATRAP peaks (with at least 50% overlap of either feature length).

### ATRAP peaks co-map with early OK-seq initiation zones

We used the OK-seq technique^38^ to map initiation zones (IZs), referred hereafter as constitutive OK-origins, in untreated JEFF cells and compared the localization of those origins to that of ATRAP peaks. We identified 7963 constitutive OK-origins across the whole genome, 5656 lying in early replicating domains (Fig. 2C and S2C, right panel). Remarkably, 80.4 % of the latter co-map with a peak and, conversely, 72.3 % of peaks co-map with an early constitutive OK-origin (Fig. 2C, E). Because of the stringent threshold of significance chosen in both techniques (Methods), some weak constitutive OK-origins and peaks within regions with high local background escaped detection. The overlap between ATRAP peaks and early constitutive origins is thus even better than mentioned above. These results validate the stressline slope showing that ATRi strongly reduces the efficiency of active compensation, and agree with previous reports showing that dormant and constitutive origins distribute in different origin classes^39–41^. In addition, genome-wide mapping of ATRAP peaks, called ATRAP-seq, provides a new method to map constitutive origins along early domains.

### ETAA1 is not involved in the control of active compensation

The role of ATR in activation of dormant origins being well established, we analyzed the impact of the main ATR activators^42^, ETAA1 and TopBP1, on the efficiency of active compensation. We first focused on ETAA1, and verified that its depletion (Fig. S3A, left panel) does not impair the cell distribution in the cycle (Fig. S3A, middle panel). Molecular combing showed that, compared to cells transfected with siCtrl, the start-point displays similar median fork speed but increased median IOD. The linearity was maintained and the slope (43) was not significantly different from that of Ref-L (F(1, 22) = 0.73, p = 0.40) (Fig. S3A, right panel), which shows that ETAA1 is not essential to reach the loose level of DRC activity required to trigger firing of dormant origins.

### TopBP1 represses dormant origin activation

Taking into account a possible impact of TopBP1 depletion on cell viability and genome stability, we determined the kinetics of TopBP1 depletion following siRNA transfection in JEFF lymphoblasts and analyzed short-term outcomes of this depletion. We found that depletion is effective, although incomplete, 24 h post-transfection and persists up to 72 h (Fig. 3A, left panels). During this period, the doubling-time of the cells was not affected (Fig. 3A, right panel). To more precisely analyze cell cycle progression under TopBP1 depletion, cells were treated as described in Fig. 3B (upper panel) and analyzed by FACS at different time points after BrdU chase. Comparison of cells transfected with siCtrl or siTopBP1 did not show differences in cell cycle progression up to 48 h post-transfection (Fig. 3B, lower panels). We conclude that TopBP1 is not essential, or not limiting, for pre-IC setting. Note that the same situation has been observed in U2OS cells depleted of MTBP^43^.

**Figure 3:**
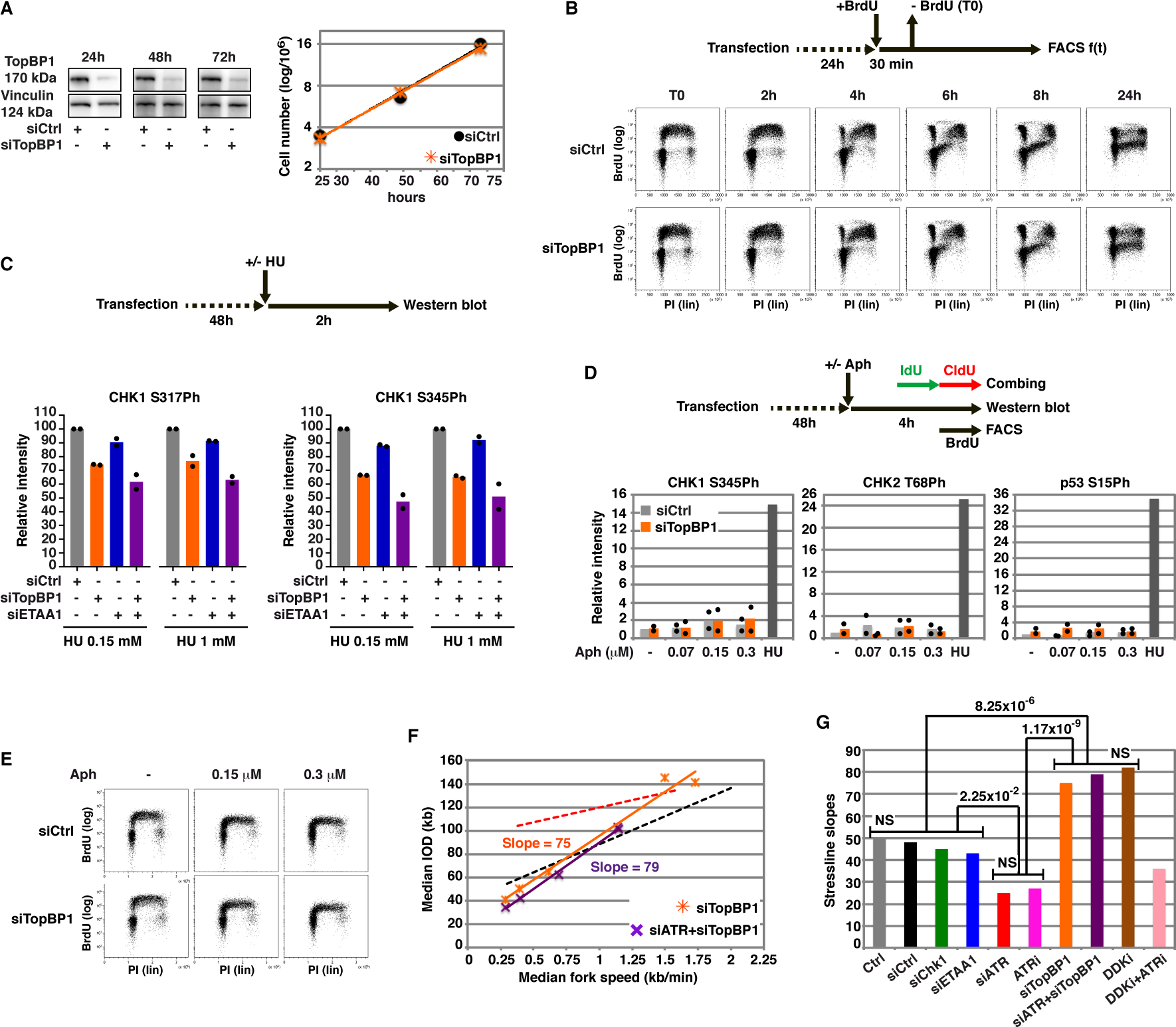
TopBP1 inhibits active compensation. **A:** Left; Western blot analysis of TopBP1 depletion at the indicated times after transfection. Right; growth curves of cells depleted or not of TopBP1. B: Upper panel; experimental procedure. Lower panels; FACS analyses comparing cell cycle progression of TopBP1 proficient and deficient cells up to 24 h after BrdU chase. C: Upper panel; experimental procedure. Lower panel; Comparison of CHK1 phosphorylation at serins 317 and 345 in cells transfected with the indicated siRNAs and treated with the indicated HU concentrations. Western blots (see Fig. S3B for an example) were performed with total extracts. For each CHK1 position, results were expressed as % of the values obtained with cells transfected with siCtrl and treated with the corresponding HU concentration. Two biologically independent experiments were performed. D: Upper panel; experimental procedure. Lower panel; Quantification of western blot (see Fig. S3C for an example) performed with total extracts of cells transfected as indicated and treated with the indicated Aph concentrations. Studied CHK1, CHK2 and p53 phospho-sites are indicated. Two biologically independent experiments were performed. Positive control: non-transfected cells treated for 2 h with HU 1mM. Normalization was done relative to the amount of non-phosphorylated protein for CHK1 and CHK2, and to ponceau staining for p53 (see Fig. S3C), band intensity is expressed relative to the value of siCtrl (-Aph) arbitrarily considered as 1. E: Cells treated as in D were analyzed by FACS. The experiment has been done once. F: Stresslines obtained with cells treated as in D, depletions are indicated (n>150 measures for each parameter). Dotted lines show stresslines corresponding to transfection with siATR (red) and siCtrl (black) as in Fig. 1H. G: Genotypes and corresponding stressline slopes are indicated. Statistical analysis (Methods) identifies three classes of proteins relative to compensation regulation: uninvolved (CHK1, ETAA1), required (ATR) and inhibitory (TopBP1).

As low doses of Aph only trigger loose DRC activation, we assessed the impact of TopBP1 depletion on CHK1 phosphorylation by treating the cells with HU 0.15mM or 1mM (Fig. 3C, top panel). Both concentrations of HU induced CHK1 phosphorylation, yet at different levels (Fig. S3B, left panel). To facilitate comparisons, we expressed the results as a percentage of the value found at each HU concentration in cells transfected with siCtrl (Fig. S3B, right panel). For each of the depletions, the relative levels of CHK1 phosphorylation proved to be similar at both HU concentrations (Supplementary Table 1A), which permitted us to pool the data (Fig. S3B, right panel). Results showed that depletion of TopBP1 significantly reduces the relative levels of CHK1 phosphorylation by some 25% at S317 and 35% at S345 (Fig. S3B, right panel and Supplemental Table 1B), suggesting redundancies in the process leading to CHK1 activation. To clarify this point, the cells were depleted of ETAA1, alone or in combination with TopBP1, and treated with HU as above. Compared to control cells, cells depleted of ETAA1 alone displayed a significant reduction of CHK1 phosphorylation at both positions, although less pronounced than in cells depleted of TopBP1 alone. Compared to single TopBP1 depletion, dual TopBP1+ETAA1 depletion significantly reduced the relative levels of CHK1 phosphorylation on S345 and S317 (Fig. 3C, lower panel and Fig. S3B and Supplemental Table 1B). Together these results show that the two ATR activators are, at least in part, functionally redundant but that other proteins, that remain to identify, also contribute to CHK1 activation. The checkpoint being modestly affected in cells depleted of TopBP1 alone, we studied the phosphorylation status of CHK1 S345, CHK2 T68 and p53 S15 in cells treated as described in Fig. 3D (upper panel). Whether or not the cells were treated with Aph (Fig. 3D, lower panel, and S3C), TopBP1 depletion impacted none of these markers, nor did we observe differences when so-treated cells were analyzed by FACS (Fig. 3E). We conclude that, with or without Aph treatment, short-term TopBP1 depletion does not promote DNA damages.

The time-frame used in previous DNA combing experiments was thus again applied to determine whether TopBP1 depletion impacts compensation (Fig. 3D, upper panel). We found that the start-point is reminiscent of that found in ATR deficient cells, and that the linearity still persists in this genetic context. Strikingly, the stressline slope (75) was significantly sharper than that of Ref-L (F(1, 21) = 12.82, p = 1.76−10-3) (Fig. 3F). TopBP1 therefore represses dormant origin activation, which opposes what would be expected from its ATR activating function and agrees with the modest impact of TopBP1 depletion on the DRC response (Fig. 3C, lower panel and Fig. S3B and Table1). In contrast, TopBP1 function at the pre-IC level, where it caps Treslin-MTBP and the GINS^12,25^, well explains this result. Indeed, a recent report has shown that the interaction of TopBP1 with the GINS precludes pol ε binding^11^, thus prevents the switch from pre-IC to CMGE. We propose that, upon stress, ATR promotes TopBP1 eviction from dormant pre-ICs, which commits those pre-ICs to firing.

### *TopBP1* is epistatic over *ATR* for active compensation

The model proposed above predicts that TopBP1 depletion should suppress ATR requirement for efficient activation of dormant origins. We thus analyzed the phenotype of cells co-depleted of TopBP1 and ATR (Fig. S3D, left panel). FACS analysis showed that dual depletion does not interfere with cell distribution in the cycle whether or not cells are treated with Aph (Fig. S3D right panel). Results obtained by molecular combing (Fig. 3F) showed that co-depletion of ATR and TopBP1 does not affect the linearity while the start-point displayed drastic fork slowing and IOD shortening, which again confirms that the linearity is independent of both the DRC and the start-point. Importantly, the slope of the stressline (79) was sharpened in the same proportion as in cells depleted of TopBP1 alone (Fig. 3G), confirming that TopBP1 depletion suppresses ATR requirement. Therefore, in contrast to a previous model that relies on the checkpoint function of TopBP1^44^, our results strongly suggest that dormant origin activation relies on the function of TopBP1 at the pre-IC level. Together, we identified three categories of proteins: uninvolved (CHK1 and ETAA1), activating (ATR) and inhibiting (TopBP1) (Fig. 3G).

### DDK down-regulates active compensation

A recent work has shown that DDK activity strengthens/stabilizes the interaction of TopBP1 with Treslin-MTBP^45^. According to our model, DDK should thus exert an inhibitory effect on active compensation. The authors used three different DDK inhibitors; TAK-931, XL413, and PHA-767491 that all reduced TopBP1/pre-IC interaction. The latter being the most efficient, we chose it for further experiments. DDK-specific phosphorylation of MCM2 S40 and S53, and CDK-specific MCM2 S27 phosphorylation^46^ were checked in cells treated with 6 μM of PHA-767491 (DDKi).

Phosphorylations of MCM2 S40 and S53 were strongly reduced while that of MCM2 S27 was not (Fig. 4A, left panel and S4A). FACS analysis showed that so-treated cells failed to enter S phase but those that have passed the G1/S transition progress through the rest of the cycle (Fig. 4A, right panel). A more detailed analysis of cell distribution in the cycle was performed by analyzing cells treated with DDKi for different periods of time then pulse-labeled with BrdU prior to FACS analysis (Fig. 4B). Results showed that DDKi modestly alter cell distribution in the cycle up to 4 h while more prolonged treatments increasingly distorted it. We therefore treated the cells for 4 h in the following experiments (Fig. 4C, upper panel). Note that such short treatment should also reduce potential consequences of DDKi off-target inhibition of CDK9^47^, a global transcription regulator.

**Figure 4:**
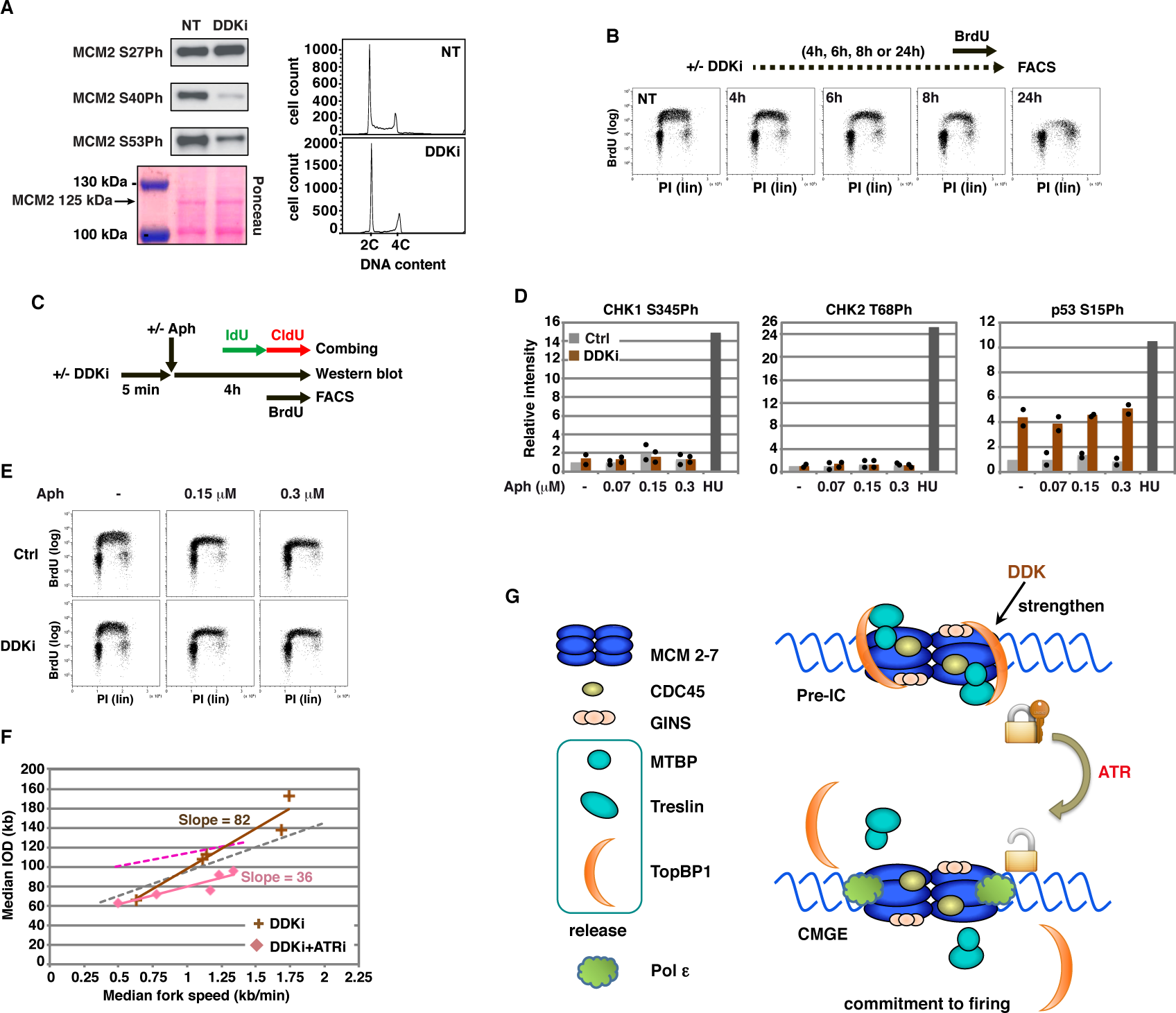
DDK inhibits compensation in lymphoblasts. **A:** Analysis of exponentially growing treated or not for 16 h with DDKi 6 μM. Left panel; Western blot analysis of MCM2 phosphorylation on the indicated serins in total cell extracts. Ponceau staining is shown as loading control. Right panel: FACS analysis of cell distribution in the cell cycle. B: Upper panel; Experimental scheme used for time-course analysis of cell distribution in the cycle in the presence of DDKi 6 μM. Lower panels; FACS data. The experiment has been done once. C: Experimental procedure relative to D, E and F. D: Western blot analysis of total extracts of cells treated as in C (western blot). Stress-induced phosphorylation of CHK1, CHK2 and p53 was quantified and expressed as in Fig. 3D. Two independent biological experiments have been performed (see Fig. S4B for an example of western blot). E: FACS analysis of cell cycle progression of cells treated as described in C (FACS). The experiment has been done once. F: Stresslines obtained with cells treated as in C (IdU and CldU labelling) with the indicated inhibitors. Dotted stresslines: as in Fig. 1H and 4F. G: Model showing how ATR and DDK cooperate to control the conversion of pre-IC into CMGE.

We analyzed the status of ATR and ATM signaling in cells treated as described in Fig. 4C. We found that, regardless of the conditions, neither CHK1 nor CHK2 showed classical activation marks. In contrast, the level of p53 and its phosphorylation on S15 increased strongly under DDKi treatment, independently of Aph (Fig. 4D and S4B). These results suggest that DDKi does not induce DNA damages under the conditions we used, but triggers the so-called licensing checkpoint^48,49^, a physiological signaling pathway that blocks normal or close-to-normal cells at the G1/S transition in response to insufficient pre-RC load. If true, DDKi offers an ideal opportunity to focus on S phase cells in which a non-limiting number of pre-RCs, and most probably of pre-ICs, have been built^48^. In support of this idea, we found that 4 h of DDKi treatment does not impact S phase progression of cells co-treated with Aph (Fig. 4E).

Molecular combing analysis showed, as previously reported^50^, that DDK inhibition modestly reduces fork speed and increases the IODs at the start-point. Strikingly, the stressline slope (82) (Fig. 4F) was not significantly different from that of TopBP1 depleted cells (F(1, 6) = 0.24, p = 0.64), which assigns DDK to the category of inhibitory proteins (Fig. 3G). Therefore, neither the role of DDK in DRC activation^50,51^ nor that in pre-RC and pre-IC building are responsible for the compensation phenotype. Instead, the stressline slope is well explained by DDK-dependent tightening of TopBP1 binding to the pre-IC.

We also analyzed cells treated with both DDKi+ATRi. Results of FACS analysis showed no difference compared to cells grown without inhibitor or with DDKi alone (Fig S4C). Molecular combing analysis revealed that dual inhibition has a strong impact on the start-point that displays a level of fork slowing resembling that caused by ATRi alone but a drastic shortening of the median IODs (Fig. 4F). These observations are consistent with previous results obtained with MCF10A cells co-treated with two other inhibitors (AZD6738 for ATR and XL413 for DDK)^50^. In striking contrast to the slope resulting from dual ATR-TopBP1 depletion, here the slope (36) was intermediate between that of control cells (50) and that of ATRi-treated (27) cells (Fig. 4E), so that it cannot be assigned to any of the previously described categories (Fig. 3G). Together results thus strongly suggest a model in which ATR and DDK exert opposing effects on TopBP1-mediated pre-IC locking, independently of each other (Fig. 4G).

## Discussion

We have developed here two novel procedures. One, called ATRAP-seq, is adapted from the Repli-seq technique^36^ and offers a robust and easy-to-implement way to map constitutive origins along early replicating domains. The other one, called stressline, relies on molecular combing analysis of how a gradient of fork slowing impacts the IODs. This procedure allowed us to show that two mechanisms are involved in compensation. First, a checkpoint-independent mechanism dictates the linear relationship linking the two parameters. Second, a checkpoint-dependent mechanism, to which the slope of the line gives access. This slope reveals the intrinsic ability of cells with different genetic backgrounds to convert pre-ICs into CMGEs. Together results obtained with this technique strongly suggest a model in which TopBP1 locks dormant origin building at the pre-IC stage. Upon replication stress, ATR promotes TopBP1 release while DDK reinforces the block. Noticeably, our model proposing that pre-ICs, not pre-RCs, are the key targets of DRC-dependent regulation of compensation does not call into question previous results showing that partial depletion of ORC or MCM subunits that makes pre-RCs limiting, and pre-ICs in consequence, reduces the compensation efficiency^18^.

Our results converge with those of previous reports to show that TopBP1 is not essential for pre-IC building in mammalian models^28–31^, which contrasts with the situation in budding and fission yeasts^25^ and in xenopus egg extracts^11,26,27^. The fact that eviction of TopBP1 from the pre-IC is mandatory for CMGE emergence^11^ thus raises the questions of what this protein is used for and why it has been conserved from yeasts to mammals. The model presented above suggests that its role has evolved towards modulation of initiation density according to the level of replication stress, a major regulatory function. The dual function of the protein, at the fork as an ATR activator and at the pre-IC as a lock to GMGE emergence, in addition suggests a regulatory loop in which eviction of TopBP1 from the pre-ICs fuels its recruitment to slowed forks, favoring amplification of DRC signaling. Note that redistribution of the Treslin-MTBP complex has been recently shown to contribute to another regulatory loop that couples replication completion to the onset of G2 phase^24^. Together, these results highlight the importance of unessential pre-IC components in S phase fitness, which explains their key role in the maintenance of genome stability.

It remains to understand how early constitutive origins escape repression since ATR activation is not likely to precede the burst of initiation that marks S phase onset. On the other hand, it is well established that DDK activity is low in G1 phase, increases around the G1/S transition and culminates in S- and G2 phases^52^. We propose that DDK is limiting at the S phase onset, and preferentially targeted to pre-RCs where it is essential for pre-IC building. In early S phase, the balance between the two kinases would thus resemble that in cells co-treated with ATRi and DDKi, in which pre-IC unlocking becomes partially independent of ATR activity.

---

**Figure S1: Compensation efficiency relies on ATR-but not CHK1-activation.**

**A:** Raw images of a typical microscopic field. Upper panel; tricolor painting of DNA fibers as in Figure 1A. lower panel; same image showing DNA counterstaining alone. **B:** Fork speed gradient resulting from treatments with the indicated Aph-concentrations. Fibers typical of the mean fork speed for each concentration are shown. **C:** Cells treated as indicated (upper panel) were analyzed by FACS (lower panel) following transfection with the indicated siRNAs. Each experiment has been done once. **D:** Western blot analysis showing how ATRi affects CHK1 phosphorylation on the indicated serins in response to treatment with a high Aph concentration. **E:** Cells treated as indicated (upper panel) were analyzed by FACS as in C (lower panel). Each experiment has been done once.

**Figure S2: Aph plus ATRi specifically reveals constitutive origins.**

**A:** Normalized density profiles of G1/S1 fractions of Repli-seq experiments of untreated cells and of cells treated with Aph or ATRi alone along the genomic region shown in Fig. 2B, showing very different patterns compared to the profiles of cells treated with ATR plus Aph (average track on top). The normalized Repli-seq signals are shown in 50kb sliding windows (with 1kb step) for visualization and range between 0 and 11 for all tracks. **B:** Normalized density profiles of G1/S1 fraction of Repli-seq experiments as in A, but along the genomic region shown in Fig. 2C. **C:** The mean normalized Repli-seq read density of G1/S1 samples with corresponding detailed heatmaps (left panels) along 0.35Mb sequences flanking the consensus peak centers across different samples. The legend of the samples and the heatmap color code are shown on the right. Each line of the heatmaps correspond to one of the consensus peaks (n=5407). The mean replication timing profiles (middle panels) and mean RFD profiles (right panels) are shown around consensus peak centers in non-treated (NT) cells. The replication timing ranges from 0 (early) to 1 (late).

**Figure S3: Impact of ETAA1 or TopBP1 depletion on compensation efficiency.**

**A:** Left panel, quality control of ETAA1 depletion 48 hours after transfection. Middle panel, cells treated as in Figure 3D (upper panel) were analyzed by FACS for cell cycle progression in the indicated growth conditions. The experiment has been done once. Right panel; ETAA1 Stressline. Dotted stresslines: as in Fig. 1H. **B:** left panel, western blot showing the phosphorylation status of CHK1 phosphorylation at S317 and S345 in cells treated as in Fig. 3C. Ponceau staining is shown as loading control. A second biologically independent experiments gave consistent results (Fig. 3C). The right panel shows the consequences of the different depletions on CHK1 phosphorylation The relative levels of CHK1 phosphorylation showing no significant differences between the two HU concentrations (T-test analysis, Supplementary Table 1A), data were pooled (HU 0.15 mM + 1 mM) to compare the outcomes of the different genotypes. (*) statistical significance at the 5% level (P-value < 0.05), (**): statistical significance at the 1% level (P-value < 0.01), (***): statistical significance at the 0.1% level (P-value < 0.001) (Supplementary Table 1B). **C:** Western blot showing the phosphorylation status of CHK1, CHK2 and p53 in cells treated as described in Fig. 3D. Growth conditions and studied phospho-sites are indicated. Ponceau staining is shown as loading control. A second biologically independent experiment gave consistent results (quantifications shown in Fig.3D). **D**: co-depletion of TopBP1 and ATR. Left panel; quality control of the depletions, vinculin is shown as loading control. Right panel; FACS analysis of the impact of dual depletion on cell cycle progression of cells treated as indicated.

**Figure S4: Impact of DDKi on cell fitness.**

**A:** Upper panel; Chromatin extracts were prepared from cells treated as indicated. Western blot shows DDKI dependent phosphorylation of MCM2 in cells treated as indicated. The experiment has been done three to four times with consistent results. Lower panel; Histogram showing signal intensities normalized relative to total MCM2 amounts. **B:** Example of western blot related to Fig. 4D. The intensity of phospho-CHK1 and phospho-CHK2 signals have been normalized relative the total amount of each protein. The intensity of p53 and p53S15 phosphorylation signals have been normalized relative to that of ponceau staining. HU (1mM) is used as positive control. **C:** Experimental scheme (upper panel). The impact of dual treatment with ATRi and DDKi on cell cycle progression was analyzed by FACS in cells treated with the indicated concentrations of Aph (lower panels).

**Table S1: Statistical analysis of ATR activator contribution to CHK1 activation.**

**A:** T-tests were conducted to assess the statistical significance of the disparity in relative intensities between cells with the same depletion(s) treated either with HU 0.15 mM or 1 mM. No significant differences were observed. Consequently, the relative intensity values for the same depletions were grouped. These grouped values were used in B for comparisons of the various depletions. **B:** T-tests were conducted to assess the statistical significance of the differences in relative intensities between cells transfected with the indicated siRNAs.

## Methods

### Cell culture and transfection

JEFF lymphoblasts were grown in RPMI 1640 + GlutaMAX-I medium (61870-010, GIBCO) supplemented with 10% foetal calf serum (CVFSVF00-0T, Eurobio) and 100 μg/mL of penicillin and streptomycin (151140-122, GIBCO) at 37 °C in a humid atmosphere containing 5% CO2. For transfections, 2−10^6^ lymphoblastoid cells were suspended in 100 μL of Nucleofector C solution (VCA-1004, Lonza Cologne AG) with 0.6 μM RNAi and transfected with the Z-001 program according to manufacturer’s instructions. Mixtures of 3 RNAi directed against ATR (HSS100876, HSS100877, HSS100878, Invitrogen), CHK1 (HSS101854, HSS101855, HSS101856, Invitrogen), TopBP1 (HSS117145, HSS117146, HSS117147, Invitrogen) or ETAA1 (HSS122911, HSS122912, HSS122913, Invitrogen) were used for transfection. The AllStars Negative Control siRNA (1027281, QIAGEN) was used for NONsi.

HCT116 cells were grown in DMEM (41965-039, GIBCO) plus 1 mM pyruvate (11360-039, GIBCO) and RPE in DMEM/F-12 (11320-074, GIBCO). Both media were supplemented with 10% foetal calf serum, 100 μg/mL of penicillin and streptomycin.

Cells treatments were performed with Aphidicolin (A0781, Sigma-Aldrich), VE-822 (2452, Axon Medchem), PHA-767491 (S2742, Selleckchem) and hydroxyurea (H8627, Sigma-Aldrich).

### FACS analyses

Cells were grown with BrdU 50 μM (5-bromo-2’-deoxyuridine, B5002, MERCK) for 30 min to label neo-synthesised DNA. 1.10^6^ cells were incubated 20 min at 37°C in 100 μL solution BD cytofix/ cytoperm (BD Pharmingen^TM^ BrdU Flow Kit, 552598, BD Bioscience), washed with PBS and re-suspended in 700 µl of staining buffer (PBS with 3% SVF). After incubation with an anti-BrdU antibody coupled to APC (Allophycocyanine), DNA was labelled with 5 µg/ml propidium iodide (IP4864-10ML, SIGMA-ALDRICH). BrdU incorporation and DNA content were measured on a FACS D Accuri Flow Cytometer C6 plus (BD Bioscience). Live cells and single cells were selected for the analysis.

### Molecular DNA combing

Molecular DNA combing was performed as previously described^32,39^. In brief, neo-synthesized DNA was labeled by two successive pulses (30 min each) with IdU 20 μM (5-iodo-2’-deoxyuridine, 57830, MERCK) and CldU 100 μM (5-chloro-2’-deoxyuridine, C6891, MERCK). Cells were embedded in agarose block and DNA was prepared and combed on silanized coverslips. IdU, CldU and DNA counterstaining were performed as previously described. Imaging was performed on an Axio Imager.Z2 microscope (Zeiss) equipped with a motorized stage to scan the slide with the Meta Imaging Series® 7.8 software (Molecular Devices). A total of 150 tracks of IdU-CldU were measured to obtain the median replication speed in kb/min. The median IOD in kb was obtained with 150 measurements. The fork density (number of forks/Mb) was measured over 200 Mb DNA.

### Protein extractions and Western blotting

Total protein extracts were prepared with 2.10^6^ cells re-suspended in 50 μL of loading buffer (B7709S, Biolabs) containing 1% SDS, 10 mM DTT and 10 mM MgCl2, then incubated with 50 U of benzonase (D0017, Millipore) for 30 min at room to digest DNA. Before electrophoresis, all samples were heated for 5 min at 95 °C. Chromatin fractionation was performed as described^53^.

Proteins were analyzed by electrophoresis in NuPAGE^TM^ 4-12% Bis-tris Gel (NP0323, INVITROGEN) or NuPAGE^TM^ 3-8% Tris-Acetate Gel (EA03755, INVITROGEN) using the running buffer MOPS SDS (NP0001, INVITROGEN) or Tris-Acetate SDS (LA0041, INVITROGEN), respectively. Proteins were transferred onto nitrocellulose membrane with an Iblot machine (kit IB301001, INVITROGEN) using program 3. Saturation was performed in PBS containing 0.05% Tween 20 and 5% BSA. Membranes were incubated with primary antibodies then secondary antibodies coupled to horseradish peroxidase overnight at 4 °C and 1 hour at room temperature, respectively. After incubation with antibodies, membranes were washed 3 times for 10 min in PBS containing 0.05% Tween 20. The chemiluminescence signals were revealed using the WesternBright ECL kit (K-12045, Advansta) and acquired with the Amersham Imager 680 (GE HealthCare). Signals on digital images were quantified with the Image Gauge software (Fujifilm).

### OK-seq data generation and processing

OK-seq libraries were generated as previously described^38^. Libraries were sequenced on an Illumina NextSeq 500 sequencing system using Paired-end (75 cycles). Data were analyzed with OKseqHMM with default parameters to compute the replication fork directionality (RFD) and determine the locations of initiation zones (IZs) genome-wide as previously described^39^.

### Repli-seq data processing

The Repli-Seq data were demultiplexed using bcl2fastq2 (v2.15.0, v2.18.12) Illumina adapters were removed using Cutadapt-1.3 (v1.3-v1.9), keeping only reads with a minimal length of ten nucleotides. The reads were mapped on the human genome (version Hg19) using bwa-mem2(v2.0pre2). The PCR duplicates were removed with the Picard (http://broadinstitute.github.io/picard, v2.25.4). Only proper paired-end reads with a minimum mapping quality score (MAPQ) of 7 were kept for downstream analysis. The read density (Dw,S1) was then computed for each 10 kb non-overlapping window (w) for each sample and normalized to the sample with the lowest number of reads. The read filtering and density were computed using the R package bamsignals v1.30.0, the read normalization and conversion to BED files with a customized script. We defined as “consensus” sample the average read density of the 6 ATRAP-seq (3 replicates of ATRAP 0.3Aph and 3 ATRAP 0.6Aph, respectively) processed as above.

### Early constitutive origins calling

Burst of replication have been mapped genome-wide on 10 kb-normalized Repli-seq G1/S1 profiles with an in-house peak caller detailly described below. For each profile, local maxima were identified by convoluting the normalized Repli-seq signal (read density) with a first derivative of a Gaussian (sigma 3 over ± 90 kb). The points of inflection were identified by convoluting the normalized Repli-seq signal with a kernel containing the second derivative of a Gaussian (sigma 3 over ± 90 kb). Peaks were called as a local maximum between two inflection points with opposite inclination. The peaks identified on the “consensus” sample called consensus peaks, and those with a mean replication timing value (S50) less than 0.4 were selected using our previous study of non-treated cells^37^ as early constitutive origins.

### Statistical analysis of stressline parallelism

To compare the stressline slopes of different conditions and groups, we performed a series of linear regression analyses of IOD and Fork Speed according to treatment, and the homogeneity of regression slopes has been assessed with the R (version 4.2.1) function anova. For each comparison, two linear regression models were built. In the first model (common slope model), we assumed a uniform relationship between the IODs and Fork Speeds across compared groups, resulting in a common slope. Conversely, the second model (vary slope model) allowed for varying slopes for the different treatment groups, aiming to investigate whether the slopes of the regression lines significantly diverged among treatment groups. We conducted an analysis of variance (anova function in R) to compare the two models and investigate whether the inclusion of interaction terms (vary slope model) significantly improved the model’s fit compared to the common slope model. The ANOVA uses a F-test to gauge the statistical significance of slope differences, as it assesses the ratio between the variance explained by the regression model and the residual variance (sum of squared residuals). A significative p-value (<0.05) suggests that the complete model (vary slopes) is a better fit for the data.

We used this analytical approach, firstly, to categorize the different treatment conditions into distinct groups/families that displayed no significant differences in their stressline slopes and, secondly, we treated these groups/families as single conditions with uniform slopes, allowing inter-group slope comparisons.

### Data availability

The raw sequencing data and the processing data will be submitted to Gene Expression Omnibus (GEO) with accession number GSEXXXXXX prior to acceptation.

### Antibodies

The following primary antibodies were used in this study:

Mouse anti-ATR (sc-515173, Santa Cruz Biotechnology)

Mouse anti-CHK1 antibody (sc-8408, Santa Cruz Biotechnology)

Rabbit anti-CHK2 (Ab109413, Abcam)

Rabbit anti-CHK1 S317Ph (12302, Cell Signaling Technology)

Rabbit anti-CHK1 S345Ph (2348, Cell Signaling Technology)

Rabbit anti-CHK2 T68Ph (2661, Cell Signaling Technology)

Mouse anti-TopBP1 (sc-271043, Santa Cruz Biotechnology)

Rabbit anti-ETAA1 (Ab122245, Abcam)

Mouse anti-Vinculin (sc-271970, Santa Cruz Biotechnology)

Mouse anti-TFIIIB90C (sc-1887, Santa Cruz Biotechnology)

Rabbit anti-MCM2 (Ab4461, Abcam)

Rabbit anti-MCM2 S27Ph (Ab109459, Abcam)

Rabbit anti-MCM2 S40Ph (Ab133243, Abcam)

Rabbit anti-MCM2 S53Ph (Ab109133, Abcam)

Mouse anti-p53 (M7001, DAKO)

Rabbit anti-p53S15Ph (9284, Cell Signaling Technology).

## Acknowledgements

The two teams contributing to the work (M.D., C-L.C.) have been supported by the Agence Nationale pour la Recherche (ANR) AAPG 2019 [19-CE12-0020-02] “TELOCHROM”. M. D.’s team is also supported by ANR AAPG 2020 [20-CE12-0027-02] “CARE-ME”. The C-L. C.’s team is also supported by ANR AAPG 2019 [19-CE12-0016-02] “ReDeFINe” and 2021 [21-CE12-0033-02] “SMART”, and by grants from the Curie Institute YPI program, the ATIP-Avenir program [ATIP/AVENIR: N◦18CT014-00] from CNRS and Plan Cancer from INSERM, and the Impulscience of Fondation Bettencourt Schueller. S.G. was supported by ATIP-Avenir and Plan Cancer for his post-doc fellowship. S.E.-H. was supported by the FRM (programme DBI20131228560). S.-J. Kim is supported by the ANR CARE-me, R.M.-G. by the ANR TELOCHROM. We acknowledge the Imaging and Cytometry Platform (UMS 3655 CNRS/US 23 INSERM) of Gustave Roussy Cancer Campus for assistance with cell sorting.

## Author contribution

M.D. conceived the project. M.D., S.K. and C.-L.C. wrote the paper. S.K. contributed to- and directed A.T. and R. M.-G. bench-work. S.-J.K. and O.B. contributed to biological experiments, including cell culture and sorting, FACS analyses and construction of Repli-Seq libraries. X.W. conducted the OK-seq experiment and Y.L. performed the OK-seq data analysis. N.A., S.G., S.E.-H. and C.-L.C. performed sequencing data and statistical analyses. F.R. provided determinant help in biological data analyses. F.R. and O.B. provided critical revision of the manuscript. S.-J. K. and N.A. contributed equally to the work.

