## Supplemental figures and table for "The balance between ATR and DDK activities controls TopBP1-mediated locking of dormant origins at the pre-IC stage"

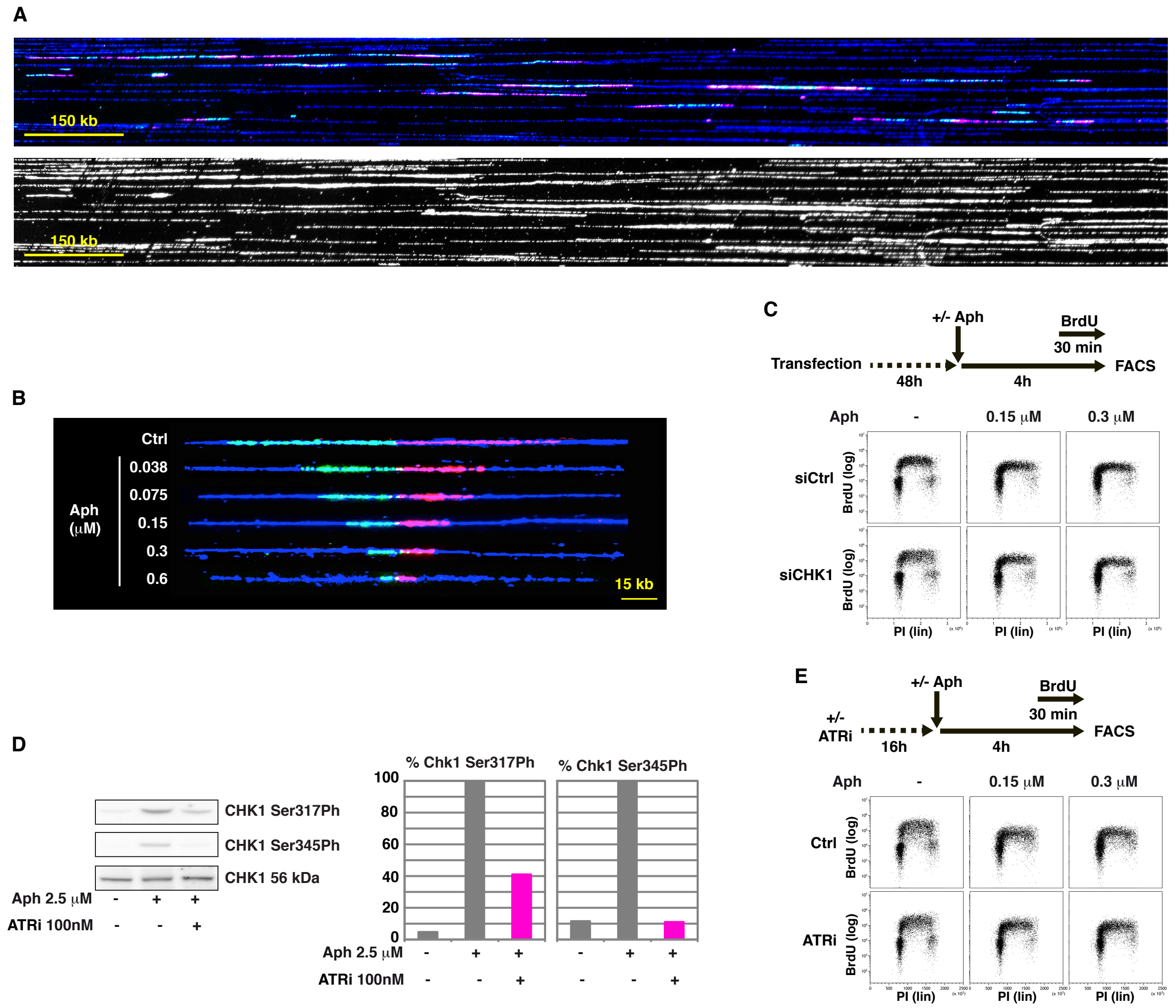


**Figure S1: Compensation efficiency relies on ATR- but not CHK1-activation.**

**A:** Raw images of a typical microscopic field. Upper panel; tricolor painting of DNA fibers as in Figure 1A. lower panel; same image showing DNA counterstaining alone. **B:** Fork speed gradient resulting from treatments with the indicated Aph-concentrations. Fibers typical of the mean fork speed for each concentration are shown. **C:** Cells treated as indicated (upper panel) were analyzed by FACS (lower panel) following transfection with the indicated siRNAs. Each experiment has been done once. **D:** Western blot analysis showing how ATRi affects CHK1 phosphorylation on the indicated serins in response to treatment with a high Aph concentration. **E:** Cells treated as indicated (upper panel) were analyzed by FACS as in C (lower panel). Each experiment has been done once.

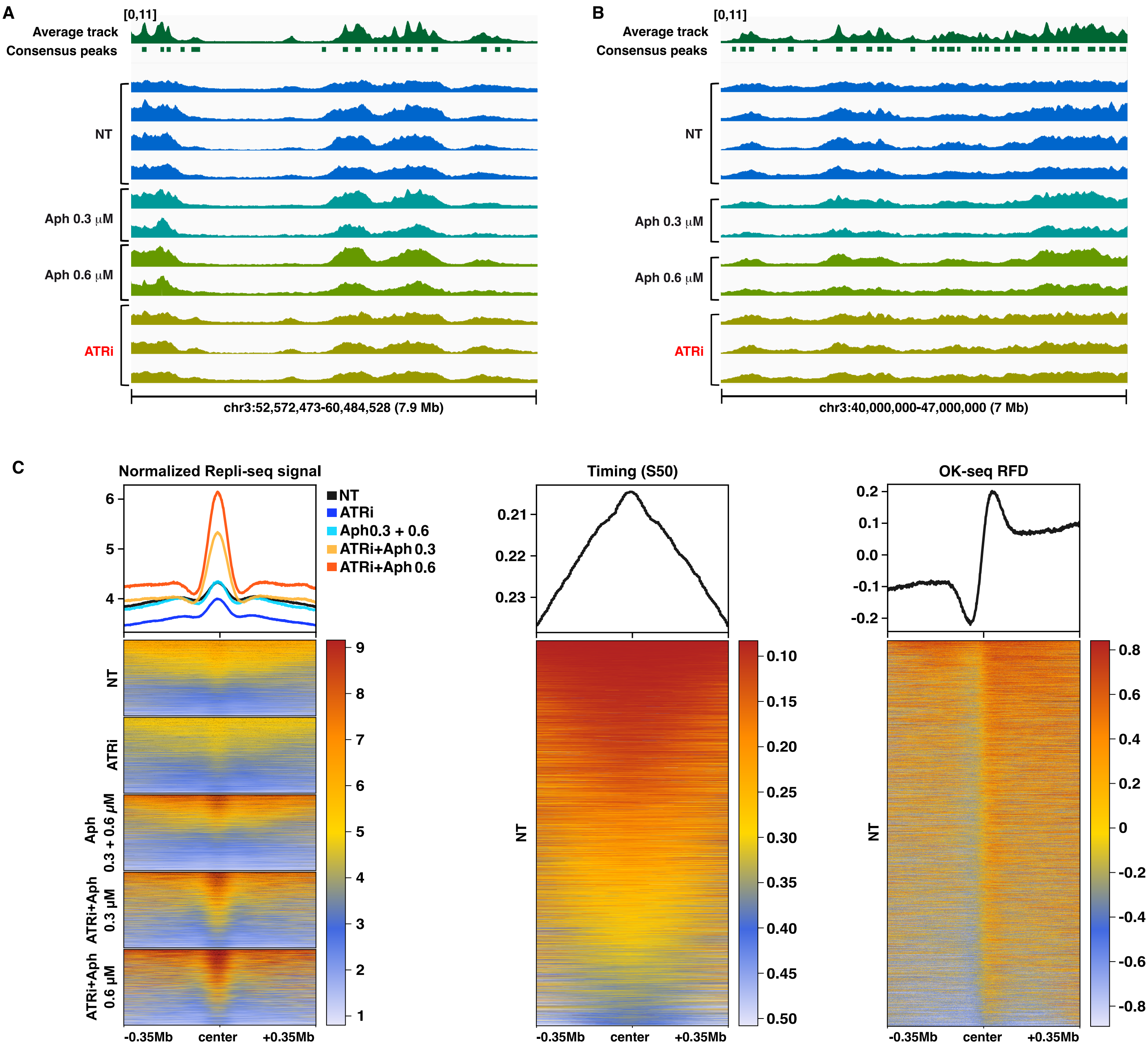


**Figure S2: Aph plus ATRi specifically reveals constitutive origins.**
**A:** Normalized density profiles of G1/S1 fractions of Repli-seq experiments of untreated cells and of cells treated with Aph or ATRi alone along the genomic region shown in Fig. 2B, showing very different patterns compared to the profiles of cells treated with ATR plus Aph (average track on top). The normalized Repli-seq signals are shown in 50kb sliding windows (with 1kb step) for visualization and range between 0 and 11 for all tracks. **B:** Normalized density profiles of G1/S1 fraction of Repli-seq experiments as in A, but along the genomic region shown in Fig. 2C. **C:** The mean normalized Repli-seq read density of G1/S1 samples with corresponding detailed heatmaps (left panels) along 0.35Mb sequences flanking the consensus peak centers across different samples. The legend of the samples and the heatmap color code are shown on the right. Each line of the heatmaps correspond to one of the consensus peaks (n=5407). The mean replication timing profiles (middle panels) and mean RFD profiles (right panels) are shown around consensus peak centers in non-treated (NT) cells. The replication timing ranges from 0 (early) to 1 (late).

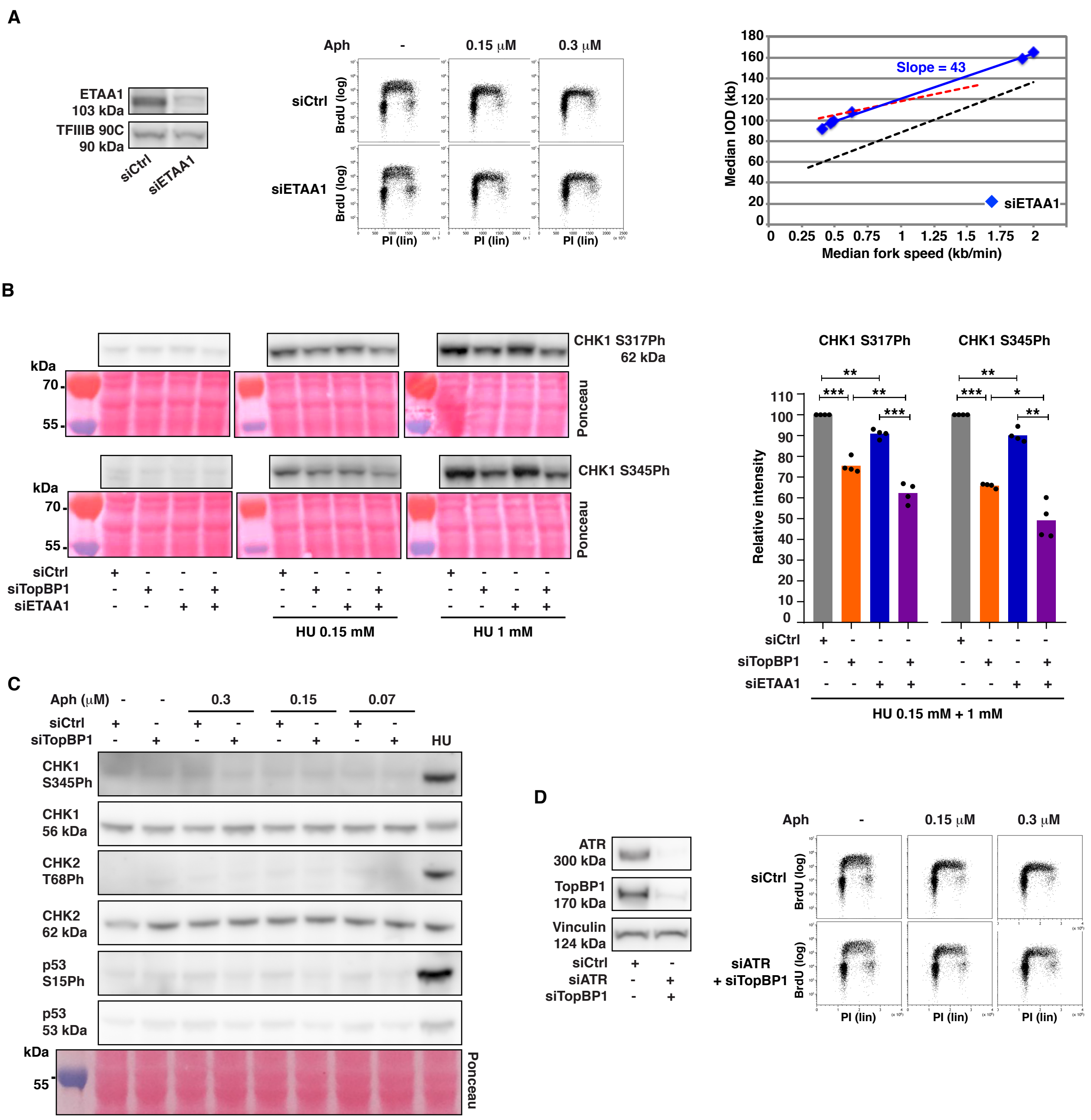


**Figure S3: Impact of ETAA1 or TopBP1 depletion on compensation efficiency.**
**A:** Left panel, quality control of ETAA1 depletion 48 hours after transfection. Middle panel, cells treated as in Figure 3D (upper panel) were analyzed by FACS for cell cycle progression in the indicated growth conditions. The experiment has been done once. Right panel; ETAA1 Stressline. Dotted stresslines: as in Fig. 1H. **B:** Left panel, western blot showing the phosphorylation status of CHK1 phosphorylation at S317 and S345 in cells treated as in Fig. 3C. Ponceau staining is shown as loading control. A second biologically independent experiments gave consistent results (Fig. 3C). The right panel shows the consequences of the different depletions on CHK1 phosphorylation The relative levels of CHK1 phosphorylation showing no significant differences between the two HU concentrations (T-test analysis, Supplementary Table 1A), data were pooled (HU 0.15 mM + 1 mM) to compare the outcomes of the different genotypes. (*) statistical significance at the 5% level (P-value < 0.05), (**): statistical significance at the 1% level (P-value < 0.01), (***): statistical significance at the 0.1% level (P-value < 0.001) (Supplementary Table 1B). **C:** Western blot showing the phosphorylation status of CHK1, CHK2 and p53 in cells treated as described in Fig. 3D. Growth conditions and studied phospho-sites are indicated. Ponceau staining is shown as loading control. A second biologically independent experiment gave consistent results (quantifications shown in Fig.3D). **D:** Co-depletion of TopBP1 and ATR. Left panel; quality control of the depletions, vinculin is shown as loading control. Right panel; FACS analysis of the impact of dual depletion on cell cycle progression of cells treated as indicated.

A

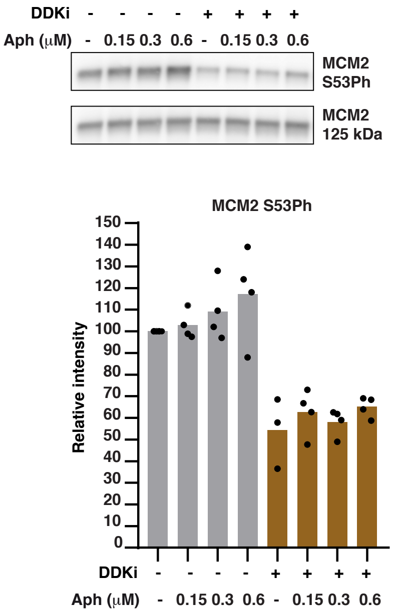


B

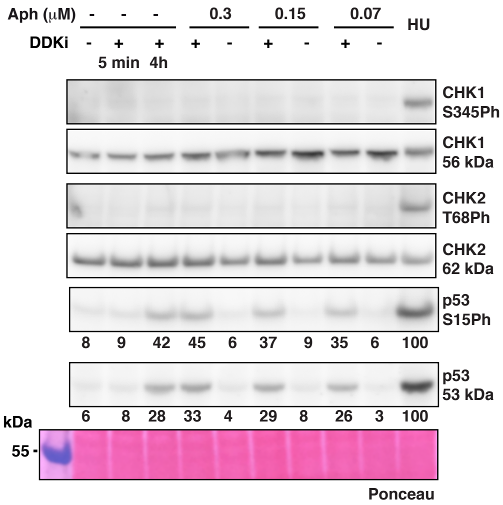


C

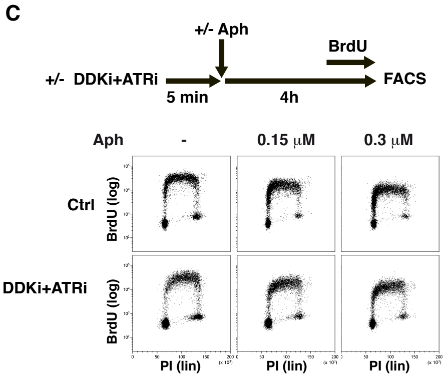


**Figure S4: Impact of DDKi on cell fitness.**
**A:** Upper panel; Chromatin extracts were prepared from cells treated as indicated. Western blot shows DDKI dependent phosphorylation of MCM2 in cells treated as indicated. The experiment has been done three to four times with consistent results. Lower panel; Histogram showing signal intensities normalized relative to total MCM2 amounts. **B:** Example of western blot related to Fig. 4D. The intensity of phospho-CHK1 and phospho-CHK2 signals have been normalized relative the total amount of each protein. The intensity of p53 and p53S15 phosphorylation signals have been normalized relative to that of ponceau staining. HU (1mM) is used as positive control. **C:** Experimental scheme (upper panel). The impact of dual treatment with ATRi and DDKi on cell cycle progression was analyzed by FACS in cells treated with the indicated concentrations of Aph (lower panels).

### Supplemental Table1

**A: Comparison HU 0.15mM vs. HU 1mM**

| | CHK1 S317Ph | CHK1 S345Ph |
|---|---|---|
| | P-value | P-value |
| siTopBP1/HU 0.15 mM **vs.** siTopBP1/HU 1 mM | 6.34 $10^{-1}$ | 3.91 $10^{-1}$ |
| siETAA1/HU 0.15 mM **vs.** siETAA1/HU 1 mM | 8.06 $10^{-1}$ | 3 $10^{-1}$ |
| siTopBP1+siETAA1/HU 0.15 mM **vs.** siTopBP1+siETAA1/HU 1 mM | 8.29 $10^{-1}$ | 7.69 $10^{-1}$ |

**B: Comparison of HU treated conditions**

| | CHK1 S317Ph | CHK1 S345Ph |
|---|---|---|
| | P-value | P-Value |
| siCtrl **vs.** siTopBP1 | 8.14 $10^{-4}$ | 8.28 $10^{-6}$ |
| siCtrl **vs.** siETAA1 | 3.44 $10^{-3}$ | 7.69 $10^{-3}$ |
| siCtlr **vs.** siTopBP1+siETAA1 | 5.67 $10^{-4}$ | 1.4 $10^{-3}$ |
| siTopBP1 **vs.** siETAA1 | 7.3 $10^{-4}$ | 2.09 $10^{-4}$ |
| siTopBP1 **vs.** siTopBP1+siETAA1 | 5.57 $10^{-3}$ | 3.13 $10^{-2}$ |
| siETAA1 **vs.** siTopBP1+siETAA1 | 3.37 $10^{-4}$ | 1.28 $10^{-3}$ |

Green: non-significant
Blue: significant (P-value < 0.05)

**Table 1: Statistical analysis of ATR activator contribution to CHK1 activation.**
**A:** T-tests were conducted to assess the statistical significance of the disparity in relative intensities between cells with the same depletion(s) treated either with HU 0.15 mM or 1 mM. No significant differences were observed. Consequently, the relative intensity values for the same depletions were grouped. These grouped values were used in B for comparisons of the various depletions. **B:** T-tests were conducted to assess the statistical significance of the differences in relative intensities between cells transfected with the indicated siRNAs.
